# The microbial metabolite Urolithin A reduces *C. difficile* toxin expression and repairs toxin-induced epithelial damage

**DOI:** 10.1101/2023.07.24.550342

**Authors:** Sweta Ghosh, Daniel Erickson, Michelle J Chua, James Collins, Venkatakrishna Rao Jala

## Abstract

*Clostridioides difficile* is a gram-positive, anaerobic, spore-forming bacterium that is responsible for antibiotic-associated pseudomembranous colitis. *Clostridioides difficile* infection (CDI) symptoms can range from diarrhea to life-threatening colon damage. Toxins produced by *C. difficile* (TcdA and TcdB) cause intestinal epithelial injury and lead to severe gut barrier dysfunction, stem cell damage, and impaired regeneration of the gut epithelium. Current treatment options for intestinal repair are limited. In this study, we demonstrate that treatment with the microbial metabolite urolithin A (UroA) attenuates CDI-induced adverse effects on the colon epithelium in a preclinical model of CDI-induced colitis. Moreover, our analysis suggests that UroA treatment protects against *C. difficile-*induced inflammation, disruption of gut barrier integrity, and intestinal tight junction proteins in the colon of CDI mice. Importantly, UroA treatment significantly reduced the expression and release of toxins from *C. difficile*, without inducing bacterial cell death. These results indicate the direct regulatory effects of UroA on bacterial gene regulation. Overall, our findings reveal a novel aspect of UroA activities, as it appears to act at both the bacterial and host levels to protect against CDI-induced colitis pathogenesis. This research sheds light on a promising avenue for the development of novel treatments for *C. difficile* infection.

**Importance:** Therapy for *C. difficile* infections includes the use of antibiotics, immunosuppressors, and fecal microbiota transplantation (FMT). However, these treatments have several drawbacks, including the loss of colonization resistance, promotion of autoimmune disorders, and the potential for unknown pathogens in donor samples. To date, the potential benefits of microbial metabolites in CDI-induced colitis have not been fully investigated. Here, we report for the first time that the microbial metabolite Urolithin A has the potential to block toxin production from *C. difficile* and enhance gut barrier function to mitigate CDI-induced colitis.

## Introduction

To cause disease, *C. difficile* must overcome colonization resistance and invade the host microbiota. Despite successful antibiotic treatment for CDI, recovery of a functional intestinal epithelial barrier in patients with CDI is impaired. The *C. difficile* toxins TcdA and TcdB are principally responsible for damage to colonic epithelial and stem cells, respectively, triggering inflammation and impairing tight junction barrier function through Rho-inhibition (1–4). Together, these can result in significant damage to the gut epithelium, leading to increased permeability, severe diarrhea, and colitis (5–7). Currently, no therapeutics are available for the treatment of *C. difficile*-induced gut barrier dysfunction or promotion of functional intestinal epithelial barrier recovery following infection. Furthermore, current antibiotic therapies disrupt the commensal microbiota, resulting in the loss of colonization resistance against *C. difficile* and altering the balance of microbial metabolites with unknown implications.

Our group and others have investigated the mechanisms and benefits of the dietary microbial metabolite urolithin A (3,8-dihydroxy-6H-benzo[c]chromen-6-one, UroA) under several pathophysiological conditions (8–14). UroA belongs to the family of urolithins and is characterized by a chemical structure containing an α-benzo-coumarin scaffold. Urolithins are produced in the gut following the microbiome-mediated transformation of the natural polyphenols ellagitannins (ETs) and ellagic acid (EA), which are present in dietary products, such as pomegranates, berries, and walnuts (15–25). Studies indicate that only 40-50% of people can produce UroA upon consuming these foods. This difference is attributed to the presence or absence of a specific bacterium or group of bacteria responsible for UroA production. When present, UroA can reach micromolar levels without toxicity in humans or mice (15, 26–30). UroA has recently been shown to reduce inflammation, enhance gut barrier function, and attenuate colitis in murine models in an aryl hydrocarbon receptor (AHR)-dependent manner (8, 26).

As *C. difficile* is affected by intestinal metabolites (e.g., bile salts) and can shape its intestinal environment by triggering inflammation, gut barrier dysfunction, and increased gut permeability, we investigated whether UroA treatment could alleviate CDI pathogenesis.

## Results

### UroA supplementation reduced CDI pathogenesis

To determine whether UroA affects CDI severity, we challenged C57BL/6J mice with 10^6^ CD2015 spores (a clinical RT027 isolate). Mice were orally administered vehicle (1% CMC, 0.1% Tween 80) or UroA (20 mg/kg) on day -6, -5, -3, -1 and daily from the day of infection. The mice were monitored for disease severity daily and euthanized on day 5 post-infection. Two of the five mice in the *C. difficile* + vehicle group died, whereas all mice (n=5) in the *C. difficile* + UroA group survived. There was a significant loss of body weight in the *C. difficile* + vehicle group compared with that in the control group (i.e., antibiotics only) (**Fig. 1A**). A standard clinical scoring system was used to evaluate the disease activity index (DAI) in infected mice. The DAI scores reflected the phenotype, where the *C. difficile* + UroA group received significantly lower scores than the vehicle group on days two and three post infection (p = 0.0146 and 0.0135, respectively; Wilcoxon rank-sum test with continuity correction, **Fig. 1B**). Shortening of the colon length in mice is a biological marker for the assessment of colonic inflammation. *C. difficile* infection caused significant shortening of colons compared to control mice. However, treatment with UroA protected the mice from *C. difficile*-induced shortening of the colon (**Fig. 1C-D**). Histopathological assessment of colon tissue showed that the UroA-treated mice had less damage and immune cell infiltration than the vehicle-treated mice (**Fig 1E**). Furthermore, analysis of inflammatory cytokines in serum suggested that UroA treatment downregulated the *C. difficile*-induced increase in IL-6, TNF-α, and IL-1β (**Fig. 1F**). Despite the amelioration of disease, the bacterial load (CFU) from the cecum/fecal samples did not show significant differences between the groups (data not shown). Mice in the UroA group lost weight compared to the control group; however, no mice succumbed to infection, and UroA significantly reduced the disease activity index and toxin-mediated inflammation.

**Figure 1.**
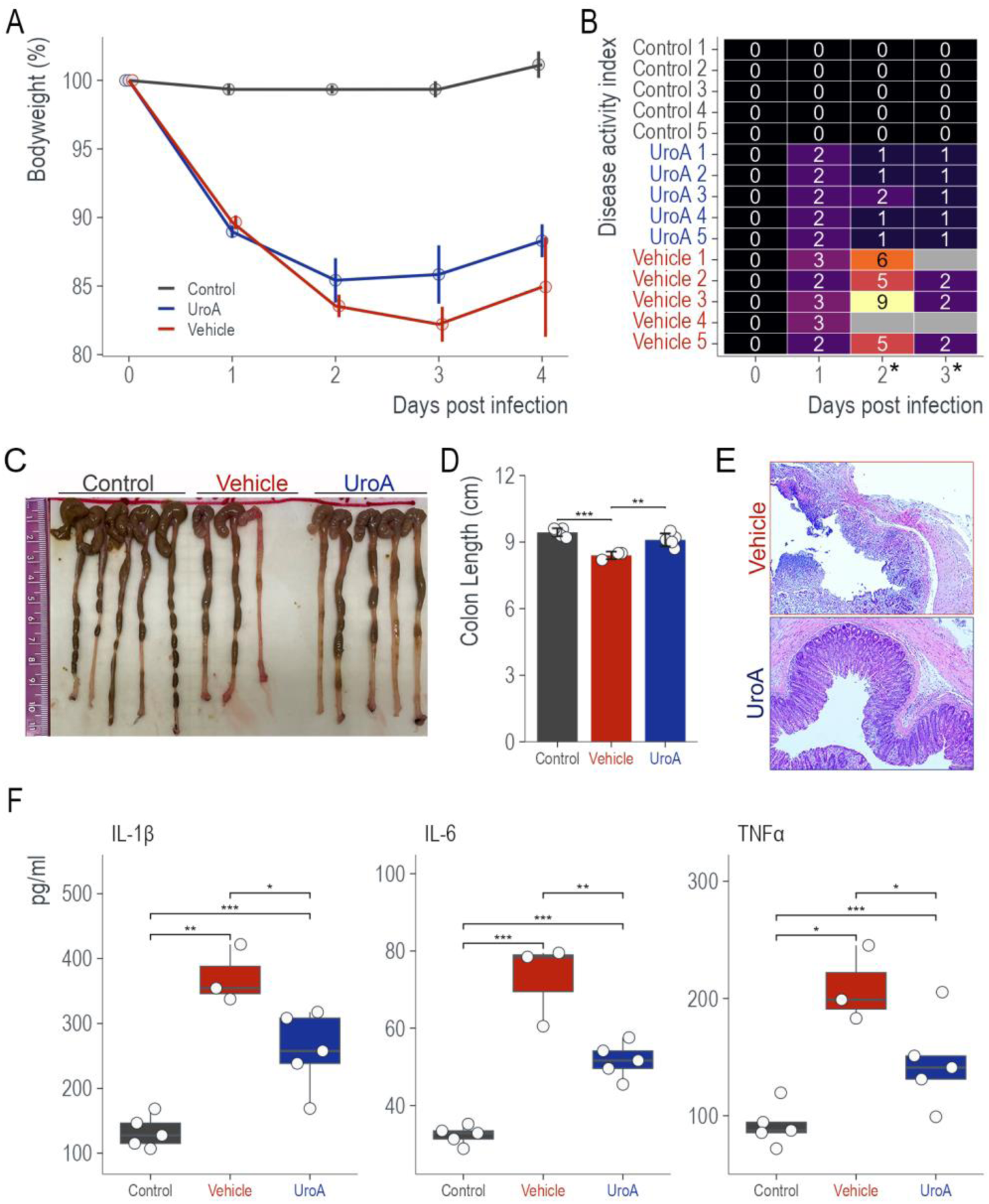
Urolithin A treatment reduces *C. difficile* pathogenesis. C57BL/6J mice (8 weeks old, n=5 per group) were subjected to *C. difficile*-induced colitis. Mice were divided into three groups: (i) control, antibiotics only, (ii) *C. difficile* + vehicle, and (iii) *C. difficile* + UroA. (**A**) Percentage body weight. Mice infected with *C. difficile* lost a significant amount of weight in both groups; however, two of the five mice (40%) in the vehicle group succumbed to infection, whereas all mice in the UroA group survived. (**B**) Disease activity index (DAI). DAI scores were given for all mice, with higher scores indicating more severe disease. The UroA group had significantly lower DAI scores on days 2 and 3 than the vehicle group (p = 0.0146 and 0.0135, respectively; Wilcoxon rank sum test with continuity correction). (**C**) Gross images of the colons. (**D**) Colon length. Both the control and UroA groups had significantly longer colons than the vehicle group (p<0.0001 and p=0.0033, respectively). (**E**) H&E staining of colonic tissue from vehicle- and uroA-treated mice following *C. difficile* infection (representative images). (**F**) Mouse serum cytokine levels of TNF-α, IL-6, and IL-1β were measured using mouse-specific ELISA. Statistics in D & F: one-way ANOVA with Tukey’s multiple comparison test.

### UroA treatment restores *C. difficile-*induced downregulation of tight junction proteins

*C. difficile* toxins can disrupt tight-junction proteins, contributing to enhanced paracellular permeability and disease pathophysiology of pseudomembranous colitis (31). Analysis of murine colons following infection showed that mice infected with *C. difficile* had significantly downregulated colon tight junction proteins (TJPs) (ZO-1, OCLN, and CLDN4). However, UroA treatment protected and restored the TJPs at the protein level (**Fig. 2A and B**), and mRNA levels (**Fig. 2C**). These results suggest that UroA treatment reduces overall inflammation and protects against CDI-induced TJP disruption.

**Figure 2.**
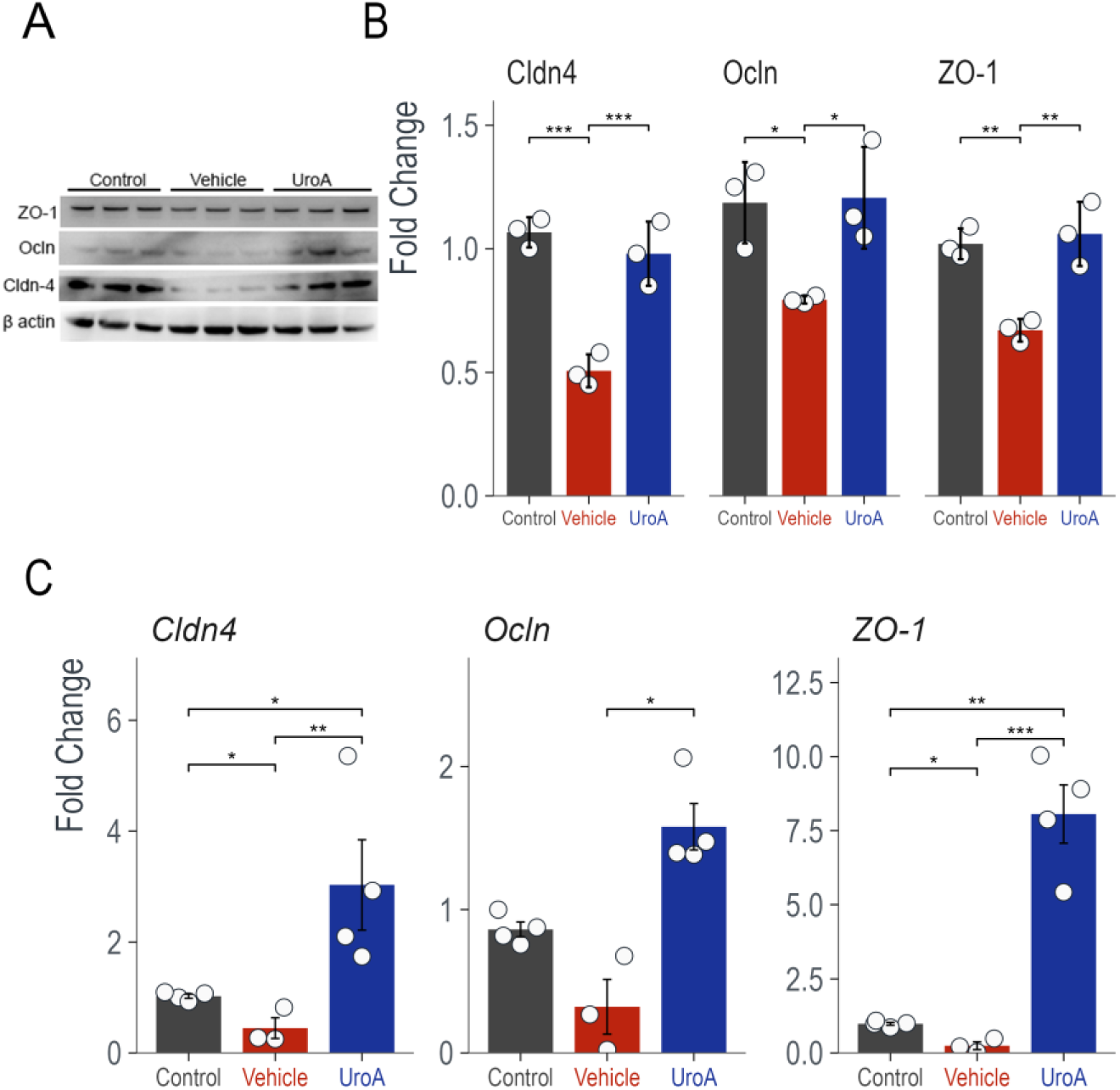
Urolithin A treatment protects CDI-induced tight junction proteins (TJPs) downregulation. *C. difficile* infection significantly reduces the TJPs claudin-4 (Cldn4), occluding (Ocln), and zonula occludens-1 (ZO-1). UroA treatment restores TJPs levels (**A**) Western blot of colonic TJPs. (**B**) Quantification of the Western blots was performed using ImageJ software. (**C**) The mRNA levels of colonic TJPs were measured using qRT-PCR with SYBR Green. Statistics are performed using one-way ANOVA with Tukey’s multiple comparison test. * p<0.05, **p<0.01, ***p<0.001

### UroA is not bactericidal

We observed no significant difference in *C. difficile* burden between the vehicle and UroA groups. This suggested that UroA does not act as an antibacterial agent against *C. difficile.* To confirm this observation and determine whether UroA elicited an antibacterial or bacteriostatic effect, we grew CD2015 in BHI medium containing increasing concentrations (0-50 µM) of UroA for 36 h and measured the OD600 every 10 min. No significant differences in doubling time or maximum OD were observed under any growth condition (**Fig. 3A**). It is possible that non-viable stationary-phase cells produce the same OD600 values as viable cells. To test whether the OD600 values masked the true bacterial viability after 36 h of growth, the samples were diluted and plated on BHI plates without the spore germinant taurocholate to enumerate the viable vegetative *C. difficile*. No significant difference was observed in CFUs for any of the UroA concentrations (p = 0.164; ANOVA, **Fig 3B**). To test for bactericidal effects more broadly, we grew the common gut commensals *Escherichia coli* and *Enterococcus faecium* in rich media containing 0-100 µM UroA. Again, no significant differences in growth or maximal OD600 were observed (p = 0.394 and p = 0.387, respectively; ANOVA; **Fig. 3C**). These data implied that UroA does not exhibit bactericidal or bacteriostatic effects at physiological concentrations.

**Figure 3.**
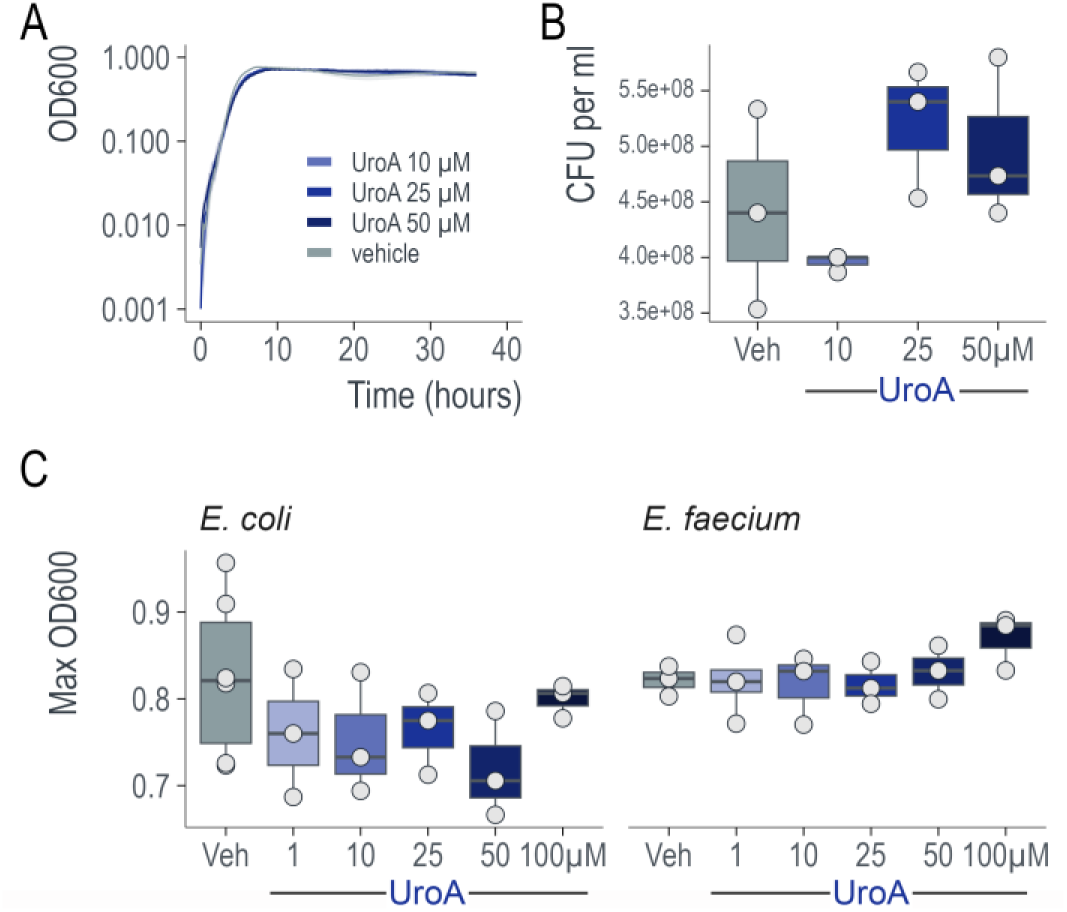
Urolithin A is neither bactericidal nor bacteriostatic. (**A**) UroA did not affect *C. difficile* growth. Three biological replicates of *C. difficile* CD2015 were grown in BHIS medium supplemented with 0-50 µM UroA. No significant differences were observed in the growth rate or maximum OD. The curves represent mean ± SD. (**B**) UroA treatment did not affect *C. difficile* viability. After 36 h of growth in BHIS + 0-50 µM UroA cells were diluted and plated onto BHIS agar plates. No significant differences were observed in the number of viable cells (ANOVA). (**C**) UroA does not affect the growth of other common intestinal microbes. *E. coli* and *E. faecium* were grow in LB and BHI media respectively containing 0-100 µM UroA. No difference in growth (max OD) was observed (ANOVA).

### UroA reduces toxin from *C. difficile* in a dose dependent manner

To determine whether UroA affects toxin expression, *C. difficile* was grown in BHI medium with or without 25 µM UroA for 24 h, and the levels of toxins in the supernatants were measured using ELISA. Upon UroA treatment, the TcdA and TcdB protein levels were significantly reduced (**Fig. 4A**). To confirm the functional reduction in toxin levels, a Vero cell rounding assay was performed with the supernatant after 24 h of growth. Cells treated with the supernatant from the *C. difficile* + vehicle group exhibited a cell-rounding phenotype, indicative of toxicity. In contrast, cells treated with the supernatant from the UroA-treated group failed to cause cell rounding (**Fig. 4B**). To determine whether the effect of UroA on *C. difficile* toxins was dose-dependent, we grew *C. difficile* in BHI medium containing increasing concentrations (0-50 µM) of UroA for 36 h and measured the toxins in the supernatant using ELISA. **Fig 4C** clearly shows the dose-dependent reduction in *C. difficile* toxin levels. These data, in combination with growth data, suggest that UroA directly interacts with *C. difficile* to downregulate toxin production in a growth-independent manner.

**Figure 4.**
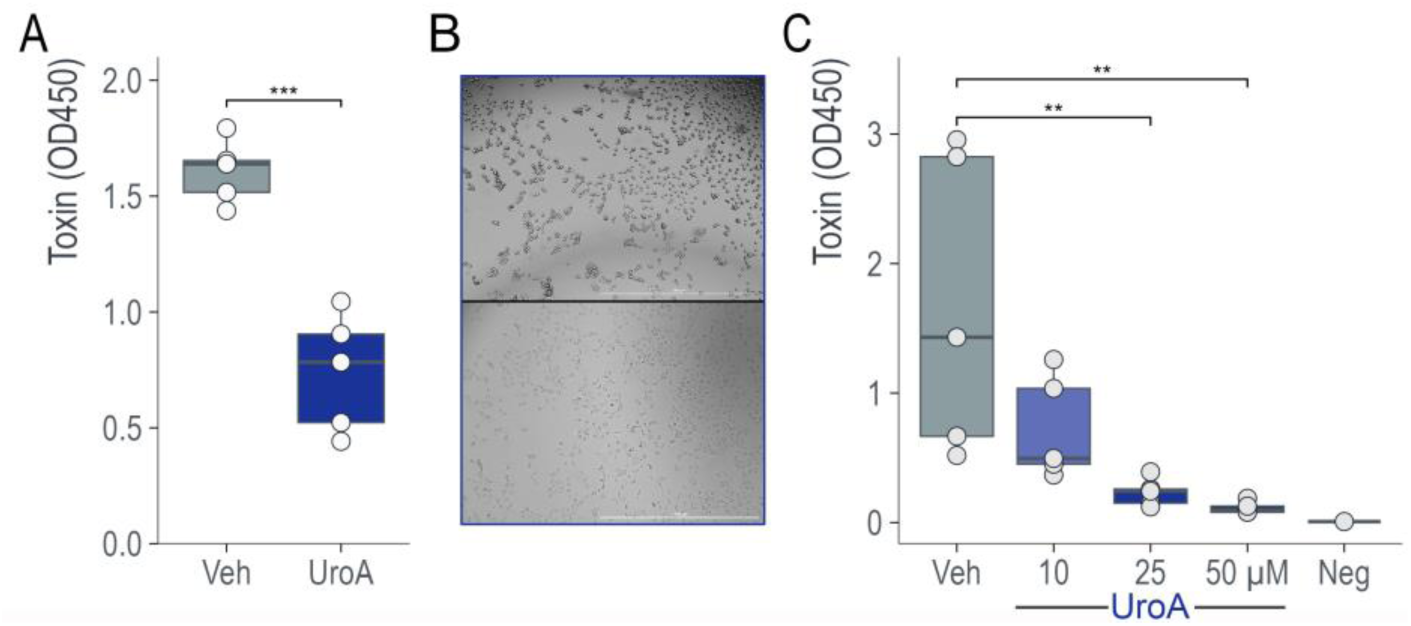
UroA reduced *C. difficile* toxin levels in a dose-dependent manner. (**A**) Five biological replicates of CD2015 were grown in BHI medium supplemented with 25 µM UroA or the vehicle control for 24 h. UroA treatment significantly reduced toxin levels (p= 0.000151, TcdA, and TcdB, as measured by non-specific ELISA). (**B**) Representative images of Vero cells after incubation with supernatant from vehicle-treated *C. difficile* (top panel) and uroA-treated *C. difficile* (bottom panel). The supernatant was collected after 24 h of growth, diluted in DMEM, and added to the Vero cells for 4 h before imaging. (**C**) Five biological replicates of CD2015 were grown in BHI medium supplemented with 10-50 µM UroA or vehicle control for 36 h. UroA treatment significantly reduced toxin levels in a dose-dependent manner, as measured by ELISA. ANOVA with post-hoc Dunnett’s test.

### UroA down regulates genes in the pathogenicity locus (PaLoc)

*C. difficile* produces toxins during stationary phase growth *in vitro*. To better understand the transcriptional landscape during this phase, we performed RNA-seq on *C. difficile* grown in the presence or absence of 25 µM UroA for 24 h. In total, 109 genes were significantly upregulated and 14 genes were downregulated in the presence of UroA (using a threshold of false discovery rate (FDR) < 0.05 Log_2_ fold-change > 1, **Fig. 5A, Table S1**). A significant portion (40%) of the upregulated genes were phage-associated from two large operons. Lysogenic phage often induces a lytic cycle when the host encounters unfavorable environmental conditions, though we did not observe any difference in viable vegetative cells. Three phosphotransferase (PTS) operons, accounting for 12 genes, were also upregulated. Several genes located in the pathogenicity locus (PaLoc) were downregulated, including *tcdA*, *tcdB*, *tcdE* (encoding a holin that mediates toxin release from *C. difficile* cells, (32, 33)), and *tcdR* (**Fig. 5B**). TcdR is an alternate sigma factor that directs transcription by recruiting RNA polymerase to the toxin gene promoters and its own promoter (34, 35). These data suggest that UroA can downregulate toxin gene expression via TcdR, either directly or through an alternative mechanism; however, further research is required to elucidate the full mechanism.

**Figure 5.**
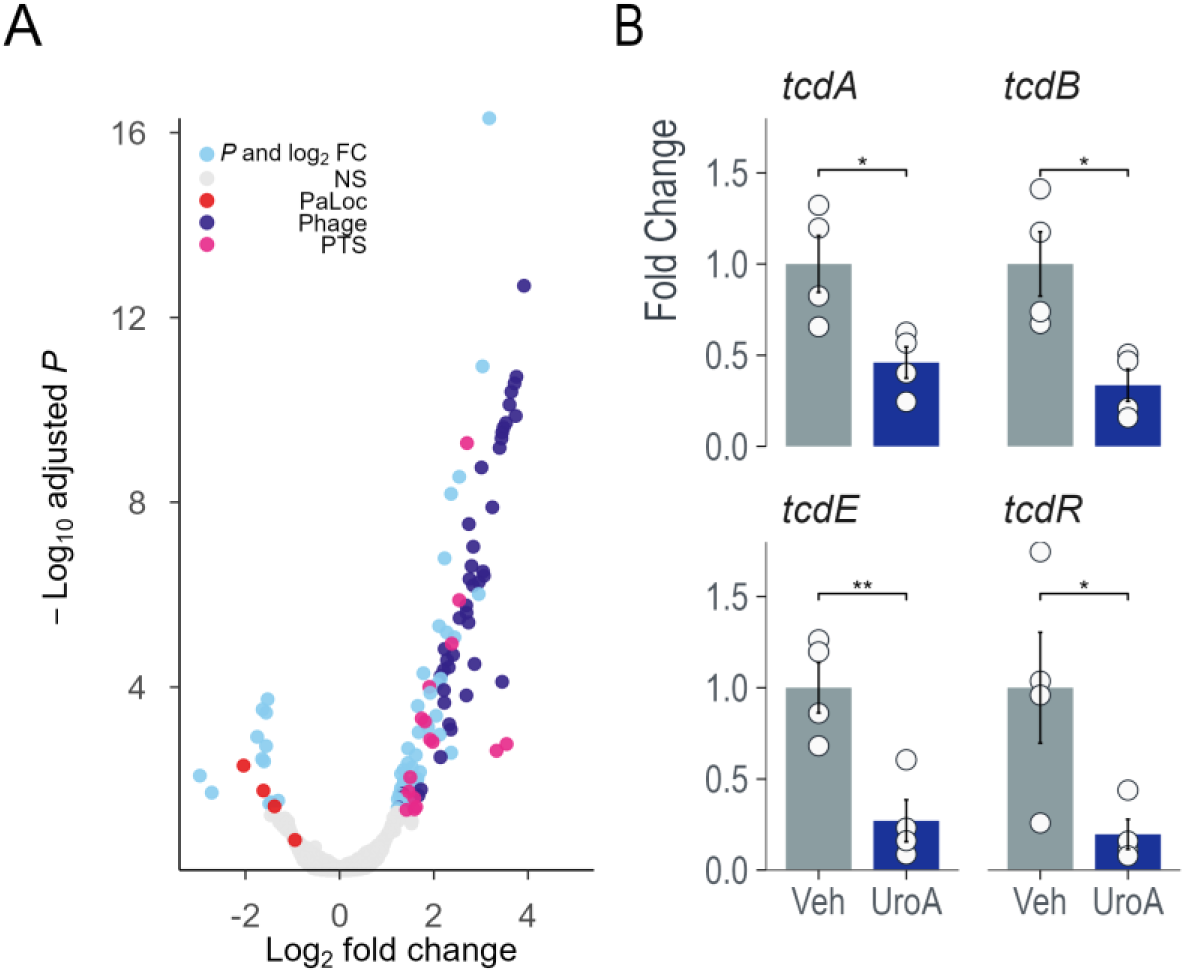
Urolithin A reduces toxin gene expression. Four biological replicates of CD2015 were grown in BHI ± 25 µM UroA or vehicle for 24 h. (**A**) RNA-seq analysis revealed that PaLoc genes were downregulated in the UroA group, whereas several PTS and phage operons were significantly upregulated. (**B**) The main toxin genes, *tcdA* and *tcdB,* were downregulated, as were *tcdE* (encoding hollin) and *tcdR* (a positive regulator of toxin gene expression).

## Discussion

During active CDI, *C. difficile* induces host pathology and inflammation, which can further exacerbate microbial dysbiosis and decrease the beneficial microbial metabolites required for gut homeostasis. Several studies have highlighted the importance of microbial metabolites in the regulation of gut barrier function by regulating both the immune and epithelial systems (36–39).

Microbial metabolites, such as short-chain fatty acids (SCFAs), indoles, purines, secondary bile acids, and polyamines, play a role in maintaining and restoring the gut barrier function and mucin production (36–39). Patients with CDI have decreased levels of tight junction proteins and increased gut barrier dysfunctions (40). Further, those patients that go on to have recurring CDI exhibit an altered gut metabolome indicative of reduced gut microbiome function, host inflammation, and reduced immune modulatory capabilities (41).

Several microbiota-derived compounds have been shown to modulate pathogen virulence factors. Of these, short chain fatty acids (SCFAs) are a well-studied class with an established role in the modulation of enteric infections by *Salmonella*, *Listeria*, *Campylobacter*, *Shigella*, and *E. coli* (42). In *C. difficile*, SCFAs have a direct inhibitory effect on growth with acetate, butyrate, propionate, and valerate, reducing the growth rate of *C. difficile* in culture (43–45). Indole, a microbial metabolite of tryptophan, reduces enterohemorrhagic *Escherichia coli O157:H7* attachment to intestinal epithelial cells and biofilm formation, and attenuates *Salmonella typhimurium* virulence and invasion, likely due to a decrease in the expression of multiple Salmonella Pathogenicity Island-1 (SPI-1) genes (46).

Toxins (TcdA and TcdB) from *C. difficile* cause epithelial damage in the intestines, leading to increased permeability and downregulation of tight junction proteins. Toxins A and B are glucosyltransferases that inactivate Rho and Ras-family GTPases within target epithelial cells, resulting in loss of structural integrity and cell death by apoptosis. Both toxins can also induce pro-inflammatory signaling pathways in the host (47, 48). A recent study by S. J. Mileto et al. (4) delineated the toxic effects of TcdA and TcdB by using specific knock-out strains. TcdA disrupts the integrity of the gut epithelial barrier, whereas TcdB damages colonic stem cells and impairs epithelial healing (4). Therefore, therapeutics that target toxin production and protect against TcdA- and TcdB-induced gut damage may mitigate CDI pathogenesis. Bezlotoxumab is an anti-toxin B monoclonal antibody that reduces the likelihood of CDI recurrence, although the mechanism by which this risk is mitigated remains unclear (49).

In the gut, UroA has been shown to reduce colon injury severity, inflammation, and intestinal permeability while improving mucosal integrity in several models (8, 26). In this study, we tested the hypothesis that supplementation with UroA protects against CDI by enhancing gut barrier function through the upregulation of tight junction proteins. Previously, our group demonstrated that treatment with UroA enhances gut barrier function, reduces inflammation, and attenuates colitis in murine models in an aryl hydrocarbon receptor (AHR)-dependent manner (8). We adopted a CDI preclinical model in which UroA supplementation significantly downregulated CDI pathogenesis (**Fig. 1**), including moderate protection from body weight loss, decreased DAI scores, reduced systemic inflammation, and mitigated CDI-induced shortening of colons. It was also evident from the H&E analysis of colon sections that UroA restored or inhibited CDI-induced gut epithelial damage (**Fig. 1**). Importantly, UroA treatment restored CDI-induced down-regulation of TJPs (**Fig. 2**).

Recent studies have highlighted that AHR activation in the intestine leads to stem cell activation, leading to the proliferation of intestinal epithelial cells (50, 51). We predict that activation of AHR by UroA may assist in the regeneration of the intestinal epithelium and induce tight junction proteins during CDI. In addition, we previously showed that UroA treatment protects the gut barrier from endotoxin- and TNF-α/IFN-γ-induced permeability and downregulation of tight junction proteins. It is also possible that UroA may reduce CDI-induced inflammatory cytokine-mediated gut barrier dysfunction. We postulate that UroA potentially regulates multiple levels to protect the host from adverse effects.

To the best of our knowledge, no studies have reported a direct effect of microbial metabolites on *C. difficile* virulence without affecting growth. A. Mahnic et al. (52) reported that bioreactors in which human derived microbiota exposed to pomegranate polyphenol extract (predominantly ellagitannins that can be converted by gut microflora to urolithin-A derivatives) in combination with clindamycin prior to *C. difficile* infection resulted in significantly less relative cytotoxicity units per *C. difficile* CFU (RCU/*C. difficile* CFU) than in the other treatments. The microbiota used was pooled from two subjects, and it is unknown whether they could produce UroA. Therefore, the mechanism of toxin reduction may have involved the production of UroA; however, this was not explicitly tested.

To determine whether UroA exhibits antibacterial activity, we grew bacteria in the presence of UroA. Our data indicated that UroA did not show any bactericidal activity. However, to our surprise, UroA treatment significantly downregulated the secretion of toxins from *C. difficile* at both protein and mRNA levels. RNA-seq revealed that UroA robustly downregulated several genes in the pathogenicity locus (PaLoc), including toxin genes and *tcdR*, a toxin gene regulator. TcdR is a central regulator of *C. difficile* toxin production. Other proteins directly regulate tcdR transcription and toxins. For example, CodY, a global transcriptional regulator, represses toxin gene expression by binding to the *tcdR* promoter region with high affinity (53, 54). The sigma factor SigD positively regulates toxin production by controlling *tcdR* transcription (55). Finally, in response to sugar availability, CcpA, a major regulator of carbon catabolite repression, binds to the promoter region or 5′ ends of several PaLoc genes, with the strongest affinity for the promoter region of tcdR (56, 57). Several PTS operons were significantly upregulated in the presence of UroA, suggesting CcpA may play a role. Further studies are required to define the UroA target and regulatory mechanisms responsible for toxin expression in *C. difficile*. In conclusion, this study provides, for the first time, an insight into how the microbial metabolite UroA can interact with the host and *C. difficile* to reduce disease severity and promote intestinal healing.

## Materials and Methods

### Bacterial strains and culture conditions

*C. difficile* CD2015 (clinical RT027 strain) was routinely grown in BHIS (BHI media (Difco) supplemented with 0.5% (w/v) yeast extract) or BHI alone for toxin assays at 37°C under anaerobic conditions (5% hydrogen, 90% nitrogen, 5% carbon dioxide). *E. coli* and *E. faecium* were grown aerobically in LB and BHI respectively.

### Mouse model of *C. difficile* infection

Mice were administered a five-antibiotic cocktail as described in J. Collins et al. (58). Briefly kanamycin (0.4 mg ml−1), gentamicin (0.035 mg ml−1), colistin (850 U ml−1), metronidazole (0.215 mg ml−1), and vancomycin (0.045 mg ml−1) were administered *ad libitum* in drinking water for 3 days. The water was switched to antibiotic-free sterile water, and 24 h later, the mice were administered an intraperitoneal injection of clindamycin (10 mg per kg (body weight)). After 24 h (day 0), the mice were challenged with 10^6^ *C. difficile* spores via oral gavage. Mice were orally administered vehicle (1% CMC, 0.1% Tween 80) or UroA (20 mg/kg) on day -6, -5, -3, -1 and daily from the day of infection. Mice were euthanized on day 5 post-infection and characterized colitis phenotype.

### Evaluation of disease activity index (DAI)

Body weight and the disease activity index (DAI) were recorded daily. The DAI was determined by combining the scores for (i) activity, (ii) posture, (iii) coat, (iv) stool consistency, and (v) eyes/nose. Each score ranged from 0 (normal) to 3 (most severe) in each area, and was scored by a blinded researcher. Body weight loss was calculated as the percentage of the difference between the original body weight (day 0) and the body weight on any day. At the end of the experiment, all mice were sacrificed and the large intestines were separated from the vermiform appendix to the anus. Colon length was measured between the cecum and proximal rectum.

### RNA-sequencing

Four independent biological replicates of *C. difficile* CD2015 were grown in BHI medium supplemented with 25 µM UroA or vehicle (DMSO) for 24h. Following growth, cells were pelleted, supernatant removed, 1 ml of RNA later added, and the pellet was stored at -80 °C until needed. Total RNA was extracted using the RNeasy kit (Qiagen), and residual DNA was removed using the TURBO DNA-free Kit (Invitrogen). RNA-Seq was performed at SeqCoast Genomics. RNA samples were subjected to ribosome depletion and sequenced on an Illumina NextSeq2000 platform using a 300-cycle flow cell kit to produce 2 × 150 bp paired reads. After demultiplexing, read trimming, and fastQC analysis, transcript expression was determined using Salmon in Python 3.10 and subsequent analysis with DESeq2 and apeglm in R v4.3.0 (59–62).

### ELISA

Toxins were measured with the Fecal *C. Difficile* Toxin AB Qualitative ELISA Assay Kit (Eagle Biosciences). Cells were removed from the supernatant via centrifugation, followed by filtration through a 0.22 µM filter. The supernatants were diluted 1:5 in PBS before following the kit instructions.

### Western blot analysis

For western blot analysis, colon tissues were homogenized and lysed using RIPA buffer containing 1X protease inhibitors (Sigma Aldrich, MO, USA), and lysates were further processed for immunoblotting as previously described (63). The membranes were probed with ZO-1, Occludin, Cldn-4, and β-actin antibodies, followed by incubation with a secondary antibody conjugated with horseradish peroxidase (Proteintech, IL, USA). The protein bands were developed with Immobilon Forte Western HRP substrate (Millipore Sigma, MA, USA) and imaged using a Bio-Rad ChemiDoc Imaging System (Hercules, CA, USA). Densitometric analysis of the bands was performed using ImageJ software (64). A list of the antibodies, sources, and dilutions used is provided in **Table S2**.

### Quantitative real-time polymerase chain reaction (qPCR)

Total RNA from colon tissues was isolated using Maxwell® 16 LEV simplyRNA tissue kits (Promega, WI, USA) following the manufacturer’s instructions. Changes in the expression of ZO-1, Occludin and Cldn4 genes were evaluated as described previously (65). Fold changes in gene expression were estimated using the ^-ΔΔ^CT method, with β-actin as a housekeeping gene control and normalized to the control.

### Measurements of serum cytokines

Mouse serum cytokine levels were measured for TNF-α, IL-6, and IL-1β using mouse-specific ELISA kits (Bio-Legend, CA, USA) following the manufacturer’s instructions.

### Histopathology and immunohistochemistry of colon tissue

Mouse colons were fixed in a 10% buffered formaldehyde solution overnight, followed by a 70% alcohol change. Fixed tissues were subjected to standard histopathological processing for paraffin embedding and 5µm paraffin sections were cut and stained with hematoxylin and eosin (H&E) by Saffron Scientific Histology Services (IL, USA).

### Data availability

Data on the normalized transcript abundance and differential analyses are shown in **Table S1**. Prior to publication of the peer-reviewed manuscript, raw data from the RNA-seq experiment shown in **Figure 5** will be available from the corresponding author upon request. Raw data will be uploaded to NCBI and made freely available upon acceptance of the peer-reviewed manuscript.

### Statistical Analysis

Statistical analysis was performed using R v4.3 (59). Details of the specific tests can be found in the relevant figure legends. *=p<0.05, **=p<0.01, ***=p<0.001. Fold change values were normalized via log2 transformation prior to statistical analysis.

## Author contributions (as outlined in CRediT author statement)

**Sweta Ghosh**: Investigation, Data Curation, Formal analysis, Writing - Review & Editing **Daniel Erickson:** Investigation, Data Curation **Michelle Chua**: Investigation, **Venkatakrishna R. Jala**: Conceptualization, Investigation, Data Curation, Formal analysis, Resources, Writing - Original Draft, Supervision, Funding acquisition, Writing - Review & Editing **James Collins**: Conceptualization, Investigation, Data Curation, Formal analysis, Resources, Writing - Original Draft, Supervision, Funding acquisition, Writing - Review & Editing.

## Funding

**JC**: This work was supported in part by a grant from the Jewish Heritage Fund for Excellence Research Recruitment Grant Program at the University of Louisville, School of Medicine, and Centers of Biomedical Research Excellence (CoBRE) Grant GM125504.

**VRJ:** VRJ is supported by NIH/NIGMS CoBRE grant (P20GM125504-01), P30ES030283 (NIH/NIEHS), NIH/NIAAA (R21AA030638), The Jewish Heritage Fund for Excellence Research Enhancement Grant, Dept. of Microbiology and Immunology and UofL-BCC.

## Declaration of interests

VRJ is a one of the co-founders of Artus Therapeutics. Other authors declare no conflicts of interest.

## REFERENCES

1. Fletcher JR, Pike CM, Parsons RJ, Rivera AJ, Foley MH, McLaren MR, Montgomery SA, Theriot CM. 2021. Clostridioides difficile exploits toxin-mediated inflammation to alter the host nutritional landscape and exclude competitors from the gut microbiota. Nat Commun 12:462.

2. Just I, Selzer J, Wilm M, von Eichel-Streiber C, Mann M, Aktories K. 1995. Glucosylation of Rho proteins by Clostridium difficile toxin B. Nature 375:500–3.

3. Hecht G, Koutsouris A, Pothoulakis C, LaMont JT, Madara JL. 1992. Clostridium difficile toxin B disrupts the barrier function of T84 monolayers. Gastroenterology 102:416–23.

4. Mileto SJ, Jarde T, Childress KO, Jensen JL, Rogers AP, Kerr G, Hutton ML, Sheedlo MJ, Bloch SC, Shupe JA, Horvay K, Flores T, Engel R, Wilkins S, McMurrick PJ, Lacy DB, Abud HE, Lyras D. 2020. Clostridioides difficile infection damages colonic stem cells via TcdB, impairing epithelial repair and recovery from disease. Proc Natl Acad Sci U S A 117:8064–8073.

5. Feltis BA, Kim AS, Kinneberg KM, Lyerly DL, Wilkins TD, Erlandsen SL, Wells CL. 1999. Clostridium difficile toxins may augment bacterial penetration of intestinal epithelium. Arch Surg 134:1235–41; discussion 1241-2.

6. Moore R, Pothoulakis C, LaMont JT, Carlson S, Madara JL. 1990. C. difficile toxin A increases intestinal permeability and induces Cl-secretion. Am J Physiol 259:G165–72.

7. Johanesen PA, Mackin KE, Hutton ML, Awad MM, Larcombe S, Amy JM, Lyras D. 2015. Disruption of the Gut Microbiome: Clostridium difficile Infection and the Threat of Antibiotic Resistance. Genes 6:1347–1360.

8. Singh R, Chandrashekharappa S, Bodduluri SR, Baby BV, Hegde B, Kotla NG, Hiwale AA, Saiyed T, Patel P, Vijay-Kumar M, Langille MGI, Douglas GM, Cheng X, Rouchka EC, Waigel SJ, Dryden GW, Alatassi H, Zhang HG, Haribabu B, Vemula PK, Jala VR. 2019. Enhancement of the gut barrier integrity by a microbial metabolite through the Nrf2 pathway. Nat Commun 10:89.

9. Ghosh S, Banerjee M, Haribabu B, Jala VR. 2022. Urolithin A attenuates arsenic-induced gut barrier dysfunction. Archives of Toxicology doi:10.1007/s00204-022-03232-2.

10. Saha P, Yeoh BS, Singh R, Chandrasekar B, Vemula PK, Haribabu B, Vijay-Kumar M, Jala VR. 2016. Gut Microbiota Conversion of Dietary Ellagic Acid into Bioactive Phytoceutical Urolithin A Inhibits Heme Peroxidases. PLoS One 11:e0156811.

11. Espin JC, Larrosa M, Garcia-Conesa MT, Tomas-Barberan F. 2013. Biological significance of urolithins, the gut microbial ellagic Acid-derived metabolites: the evidence so far. Evid Based Complement Alternat Med 2013:270418.

12. Tomas-Barberan FA, Gonzalez-Sarrias A, Garcia-Villalba R, Nunez-Sanchez MA, Selma MV, Garcia-Conesa MT, Espin JC. 2016. Urolithins, the rescue of ’old’ metabolites to understand a ’new’ concept: metabotypes as a nexus between phenolic metabolism, microbiota dysbiosis and host health status. Mol Nutr Food Res doi:10.1002/mnfr.201500901.

13. Ryu D, Mouchiroud L, Andreux PA, Katsyuba E, Moullan N, Nicolet-Dit-Felix AA, Williams EG, Jha P, Lo Sasso G, Huzard D, Aebischer P, Sandi C, Rinsch C, Auwerx J. 2016. Urolithin A induces mitophagy and prolongs lifespan in C. elegans and increases muscle function in rodents. Nat Med 22:879–88.

14. García-Villalba R, Giménez-Bastida JA, Cortés-Martín A, Ávila-Gálvez M, Tomás-Barberán FA, Selma MV, Espín JC, González-Sarrías A. 2022. Urolithins: a comprehensive update on their metabolism, bioactivity, and associated gut microbiota. Mol Nutr Food Res doi:10.1002/mnfr.202101019:e2101019.

15. Gonzalez-Sarrias A, Gimenez-Bastida JA, Garcia-Conesa MT, Gomez-Sanchez MB, Garcia-Talavera NV, Gil-Izquierdo A, Sanchez-Alvarez C, Fontana-Compiano LO, Morga-Egea JP, Pastor-Quirante FA, Martinez-Diaz F, Tomas-Barberan FA, Espin JC. 2010. Occurrence of urolithins, gut microbiota ellagic acid metabolites and proliferation markers expression response in the human prostate gland upon consumption of walnuts and pomegranate juice. Mol Nutr Food Res 54:311–22.

16. Cerda B, Periago P, Espin JC, Tomas-Barberan FA. 2005. Identification of urolithin a as a metabolite produced by human colon microflora from ellagic acid and related compounds. J Agric Food Chem 53:5571–6.

17. Cerda B, Espin JC, Parra S, Martinez P, Tomas-Barberan FA. 2004. The potent in vitro antioxidant ellagitannins from pomegranate juice are metabolised into bioavailable but poor antioxidant hydroxy-6H-dibenzopyran-6-one derivatives by the colonic microflora of healthy humans. Eur J Nutr 43:205–20.

18. Garcia-Villalba R, Beltran D, Espin JC, Selma MV, Tomas-Barberan FA. 2013. Time course production of urolithins from ellagic acid by human gut microbiota. J Agric Food Chem 61:8797–806.

19. Tomas-Barberan FA, Garcia-Villalba R, Gonzalez-Sarrias A, Selma MV, Espin JC. 2014. Ellagic acid metabolism by human gut microbiota: consistent observation of three urolithin phenotypes in intervention trials, independent of food source, age, and health status. J Agric Food Chem 62:6535–8.

20. Cerda B, Tomas-Barberan FA, Espin JC. 2005. Metabolism of antioxidant and chemopreventive ellagitannins from strawberries, raspberries, walnuts, and oak-aged wine in humans: identification of biomarkers and individual variability. J Agric Food Chem 53:227–35.

21. Seeram NP, Zhang Y, McKeever R, Henning SM, Lee RP, Suchard MA, Li Z, Chen S, Thames G, Zerlin A, Nguyen M, Wang D, Dreher M, Heber D. 2008. Pomegranate juice and extracts provide similar levels of plasma and urinary ellagitannin metabolites in human subjects. J Med Food 11:390–4.

22. Seeram NP, Henning SM, Zhang Y, Suchard M, Li Z, Heber D. 2006. Pomegranate juice ellagitannin metabolites are present in human plasma and some persist in urine for up to 48 hours. J Nutr 136:2481–5.

23. Nunez-Sanchez MA, Garcia-Villalba R, Monedero-Saiz T, Garcia-Talavera NV, Gomez-Sanchez MB, Sanchez-Alvarez C, Garcia-Albert AM, Rodriguez-Gil FJ, Ruiz-Marin M, Pastor-Quirante FA, Martinez-Diaz F, Yanez-Gascon MJ, Gonzalez-Sarrias A, Tomas-Barberan FA, Espin JC. 2014. Targeted metabolic profiling of pomegranate polyphenols and urolithins in plasma, urine and colon tissues from colorectal cancer patients. Mol Nutr Food Res 58:1199–211.

24. Cortés-Martín A, García-Villalba R, González-Sarrías A, Romo-Vaquero M, Loria-Kohen V, Ramírez-de-Molina A, Tomás-Barberán FA, Selma MV, Espín JC. 2018. The gut microbiota urolithin metabotypes revisited: the human metabolism of ellagic acid is mainly determined by aging. Food Funct 9:4100–4106.

25. Gonzalez-Sarrias A, Garcia-Villalba R, Romo-Vaquero M, Alasalvar C, Orem A, Zafrilla P, Tomas-Barberan FA, Selma MV, Espin JC. 2017. Clustering according to urolithin metabotype explains the interindividual variability in the improvement of cardiovascular risk biomarkers in overweight-obese individuals consuming pomegranate: A randomized clinical trial. Mol Nutr Food Res 61.

26. Larrosa M, Gonzalez-Sarrias A, Yanez-Gascon MJ, Selma MV, Azorin-Ortuno M, Toti S, Tomas-Barberan F, Dolara P, Espin JC. 2010. Anti-inflammatory properties of a pomegranate extract and its metabolite urolithin-A in a colitis rat model and the effect of colon inflammation on phenolic metabolism. J Nutr Biochem 21:717–25.

27. Espin JC, Gonzalez-Barrio R, Cerda B, Lopez-Bote C, Rey AI, Tomas-Barberan FA. 2007. Iberian pig as a model to clarify obscure points in the bioavailability and metabolism of ellagitannins in humans. J Agric Food Chem 55:10476–85.

28. Heilman J, Andreux P, Tran N, Rinsch C, Blanco-Bose W. 2017. Safety assessment of Urolithin A, a metabolite produced by the human gut microbiota upon dietary intake of plant derived ellagitannins and ellagic acid. Food Chem Toxicol 108:289–297.

29. Tomas-Barberan FA, Gonzalez-Sarrias A, Garcia-Villalba R, Nunez-Sanchez MA, Selma MV, Garcia-Conesa MT, Espin JC. 2017. Urolithins, the rescue of “old” metabolites to understand a “new” concept: Metabotypes as a nexus among phenolic metabolism, microbiota dysbiosis, and host health status. Mol Nutr Food Res 61.

30. Andreux PA, Blanco-Bose W, Ryu D, Burdet F, Ibberson M, Aebischer P, Auwerx J, Singh A, Rinsch C. 2019. The mitophagy activator urolithin A is safe and induces a molecular signature of improved mitochondrial and cellular health in humans. Nature Metabolism 1:595–603.

31. Nusrat A, von Eichel-Streiber C, Turner JR, Verkade P, Madara JL, Parkos CA. 2001. Clostridium difficile toxins disrupt epithelial barrier function by altering membrane microdomain localization of tight junction proteins. Infect Immun 69:1329–36.

32. Govind R, Fitzwater L, Nichols R. 2015. Observations on the Role of TcdE Isoforms in Clostridium difficile Toxin Secretion. J Bacteriol 197:2600–9.

33. Govind R, Dupuy B. 2012. Secretion of Clostridium difficile toxins A and B requires the holin-like protein TcdE. PLoS Pathog 8:e1002727.

34. Mani N, Lyras D, Barroso L, Howarth P, Wilkins T, Rood JI, Sonenshein AL, Dupuy B. 2002. Environmental response and autoregulation of Clostridium difficile TxeR, a sigma factor for toxin gene expression. J Bacteriol 184:5971–8.

35. Mani N, Dupuy B. 2001. Regulation of toxin synthesis in Clostridium difficile by an alternative RNA polymerase sigma factor. Proc Natl Acad Sci U S A 98:5844–9.

36. Sonnenburg ED, Sonnenburg JL. 2014. Starving our microbial self: the deleterious consequences of a diet deficient in microbiota-accessible carbohydrates. Cell Metab 20:779–786.

37. Bansal T, Alaniz RC, Wood TK, Jayaraman A. 2010. The bacterial signal indole increases epithelial-cell tight-junction resistance and attenuates indicators of inflammation. Proc Natl Acad Sci U S A 107:228–33.

38. Cipriani S, Mencarelli A, Chini MG, Distrutti E, Renga B, Bifulco G, Baldelli F, Donini A, Fiorucci S. 2011. The bile acid receptor GPBAR-1 (TGR5) modulates integrity of intestinal barrier and immune response to experimental colitis. PLoS One 6:e25637.

39. Wang JY, McCormack SA, Viar MJ, Johnson LR. 1991. Stimulation of proximal small intestinal mucosal growth by luminal polyamines. Am J Physiol 261:G504–11.

40. Chen S, Sun C, Wang H, Wang J. 2015. The Role of Rho GTPases in Toxicity of Clostridium difficile Toxins. Toxins (Basel) 7:5254–67.

41. Dawkins JJ, Allegretti JR, Gibson TE, McClure E, Delaney M, Bry L, Gerber GK. 2022. Gut metabolites predict Clostridioides difficile recurrence. Microbiome 10:87.

42. Sun Y, O’Riordan MX. 2013. Regulation of bacterial pathogenesis by intestinal short-chain Fatty acids. Adv Appl Microbiol 85:93–118.

43. Hryckowian AJ, Van Treuren W, Smits SA, Davis NM, Gardner JO, Bouley DM, Sonnenburg JL. 2018. Microbiota-accessible carbohydrates suppress Clostridium difficile infection in a murine model. Nat Microbiol 3:662–669.

44. McDonald JAK, Mullish BH, Pechlivanis A, Liu Z, Brignardello J, Kao D, Holmes E, Li JV, Clarke TB, Thursz MR, Marchesi JR. 2018. Inhibiting Growth of Clostridioides difficile by Restoring Valerate, Produced by the Intestinal Microbiota. Gastroenterology 155:1495–1507.e15.

45. Kondepudi KK, Ambalam P, Nilsson I, Wadström T, Ljungh A. 2012. Prebiotic-non-digestible oligosaccharides preference of probiotic bifidobacteria and antimicrobial activity against Clostridium difficile. Anaerobe 18:489–97.

46. Kohli N, Crisp Z, Riordan R, Li M, Alaniz RC, Jayaraman A. 2018. The microbiota metabolite indole inhibits Salmonella virulence: Involvement of the PhoPQ two-component system. PLoS One 13:e0190613.

47. Janoir C. 2016. Virulence factors of Clostridium difficile and their role during infection. Anaerobe 37:13–24.

48. Pechine S, Collignon A. 2016. Immune responses induced by Clostridium difficile. Anaerobe 41:68–78.

49. Villafuerte Gálvez JA, Kelly CP. 2017. Bezlotoxumab: anti-toxin B monoclonal antibody to prevent recurrence of Clostridium difficile infection. Expert Rev Gastroenterol Hepatol 11:611–622.

50. Metidji A, Omenetti S, Crotta S, Li Y, Nye E, Ross E, Li V, Maradana MR, Schiering C, Stockinger B. 2018. The Environmental Sensor AHR Protects from Inflammatory Damage by Maintaining Intestinal Stem Cell Homeostasis and Barrier Integrity. Immunity 49:353–362.e5.

51. Shah K, Maradana MR, Joaquina Delàs M, Metidji A, Graelmann F, Llorian M, Chakravarty P, Li Y, Tolaini M, Shapiro M, Kelly G, Cheshire C, Bhurta D, Bharate SB, Stockinger B. 2022. Cell-intrinsic Aryl Hydrocarbon Receptor signalling is required for the resolution of injury-induced colonic stem cells. Nature Communications 13:1827.

52. Mahnic A, Auchtung JM, Poklar Ulrih N, Britton RA, Rupnik M. 2020. Microbiota in vitro modulated with polyphenols shows decreased colonization resistance against Clostridioides difficile but can neutralize cytotoxicity. Sci Rep 10:8358.

53. Dineen SS, Villapakkam AC, Nordman JT, Sonenshein AL. 2007. Repression of Clostridium difficile toxin gene expression by CodY. Mol Microbiol 66:206–19.

54. Dineen SS, McBride SM, Sonenshein AL. 2010. Integration of metabolism and virulence by Clostridium difficile CodY. J Bacteriol 192:5350–62.

55. El Meouche I, Peltier J, Monot M, Soutourina O, Pestel-Caron M, Dupuy B, Pons JL. 2013. Characterization of the SigD regulon of C. difficile and its positive control of toxin production through the regulation of tcdR. PLoS One 8:e83748.

56. Antunes A, Camiade E, Monot M, Courtois E, Barbut F, Sernova NV, Rodionov DA, Martin-Verstraete I, Dupuy B. 2012. Global transcriptional control by glucose and carbon regulator CcpA in Clostridium difficile. Nucleic Acids Res 40:10701–18.

57. Antunes A, Martin-Verstraete I, Dupuy B. 2011. CcpA-mediated repression of Clostridium difficile toxin gene expression. Mol Microbiol 79:882–99.

58. Collins J, Auchtung JM, Schaefer L, Eaton KA, Britton RA. 2015. Humanized microbiota mice as a model of recurrent Clostridium difficile disease. Microbiome 3:35.

59. 59. R Core Team. 2023. R: A Language and Environment for Statistical Computing, R Foundation for Statistical Computing, https://www.R-project.org/.

60. Patro R, Duggal G, Love MI, Irizarry RA, Kingsford C. 2017. Salmon provides fast and bias-aware quantification of transcript expression. Nat Methods 14:417–419.

61. Love MI, Huber W, Anders S. 2014. Moderated estimation of fold change and dispersion for RNA-seq data with DESeq2. Genome Biol 15:550.

62. Zhu A, Ibrahim JG, Love MI. 2019. Heavy-tailed prior distributions for sequence count data: removing the noise and preserving large differences. Bioinformatics 35:2084–2092.

63. Ghosh S, Singh R, Vanwinkle ZM, Guo H, Vemula PK, Goel A, Haribabu B, Jala VR. 2022. Microbial metabolite restricts 5-fluorouracil-resistant colonic tumor progression by sensitizing drug transporters via regulation of FOXO3-FOXM1 axis. Theranostics 12:5574–5595.

64. Schneider CA, Rasband WS, Eliceiri KW. 2012. NIH Image to ImageJ: 25 years of image analysis. Nat Methods 9:671–5.

65. Ghosh S, Banerjee M, Haribabu B, Jala VR. 2022. Urolithin A attenuates arsenic-induced gut barrier dysfunction. Arch Toxicol doi:10.1007/s00204-022-03232-2.

